# Neuroplasticity associated with changes in conversational turn-taking following a family-based intervention

**DOI:** 10.1101/2020.10.30.362723

**Authors:** Rachel R. Romeo, Julia A. Leonard, Hannah M. Grotzinger, Sydney T. Robinson, Megumi E. Takada, Allyson P. Mackey, Ethan Scherer, Meredith L. Rowe, Martin R. West, John D. E. Gabrieli

**Affiliations:** Department of Brain and Cognitive Sciences and McGovern Institute for Brain Research, Massachusetts Institute of Technology; Center for Education Policy Research, Harvard University; Graduate School of Education, Harvard University; Integrated Learning Initiative, Massachusetts Institute of Technology

**Keywords:** Neuroplasticity, conversational turns, socioeconomic status, family-based intervention, language development, LENA

## Abstract

Children’s early language environments are associated with linguistic, cognitive, and academic development, as well as concurrent brain structure and function. This study investigated neurodevelopmental mechanisms linking language input to development by measuring neuroplasticity associated with an intervention designed to enhance language environments of families primarily from lower socioeconomic backgrounds. Families of 52 4-to-6 year-old children were randomly assigned to a 9-week, interactive, family-based intervention or no-contact control group. Children completed pre- and post-assessments of verbal and nonverbal cognition (n=52), structural magnetic resonance imaging (n=45), and home auditory recordings of language exposure (n=39). Families who completed the intervention exhibited greater increases in adult-child conversational turns, and changes in turn-taking mediated intervention effects on language and executive functioning measures. Collapsing across groups, turn-taking changes were also positively correlated with cortical thickening in left inferior frontal and supramarginal gyri, the latter of which mediated relationships between changes in turn-taking and children’s language development. This is the first study of longitudinal neuroplasticity in response to changes in children’s language environments, and findings suggest that conversational turns support language development through cortical growth in language and social processing regions. This has implications for early interventions to enhance children’s language environments to support neurocognitive development.

## Introduction

Children’s early language environments are intimately associated with their trajectories of language development, which in turn predict cognitive and academic development across childhood (Gilkerson et al., 2018; Huttenlocher et al., 2002; Rowe, 2012; Walker et al., 1994). There is immense variation in children’s language environments, and children growing up in homes with lower socioeconomic status (SES) receive, on average, quantitatively and qualitatively different language exposure than peers from higher-SES backgrounds (for review, see Pace et al., 2017; Rowe, 2018; Schwab & Lew-Williams, 2016). Further, the quantity and quality of children’s early language experience significantly mediate relationships between SES and children’s language scores (Hoff, 2003; Romeo, Leonard, et al., 2018; Romeo, Segaran, et al., 2018; Rowe & Goldin-Meadow, 2009), indicating experiential mechanisms which contribute to SES differences in language development.

Neuroimaging studies have revealed structural and functional neural mechanisms relating children’s language input to their language skills. A study of 4-to-6 year-old children found that the frequency of adult-child conversational turns was positively correlated with brain activation during language processing in left inferior frontal regions (Romeo, Leonard, et al., 2018), as well as the strength of white matter connectivity in the left arcuate and superior longitudinal fasciculi (Romeo, Segaran, et al., 2018), both of which mediated relationships between conversational turns and SES-related differences in language. Similarly, in a study of 5-to-9 year-old children, the quantity of adult words and adult-child conversational turns correlated with the surface area of left perisylvian cortical regions, which mediated SES relationships with children’s reading skills (Merz et al., 2020). Additionally, in infants from lower-SES homes, the quantity of language input in the first year of life is related to neuro-oscillatory patterns across multiple brain regions (Brito et al., 2020; Pierce et al., 2020). Together, these studies suggest strong relationships between children’s language experience and brain measures from infancy through childhood; however, these studies have all employed cross-sectional, correlational methods. Longitudinal controlled intervention studies are necessary to make causal claims about the role of language input on brain development.

Despite numerous studies of children’s neuroplasticity resulting from child-facing interventions (for review, see Weyandt, et al., 2020), fewer studies have investigated neural effects resulting from interventions specifically targeting parents or families as change agents. Clinically oriented interventions for parents of medically compromised infants, such as those born premature, find enhanced neural maturation in infants whose mothers received sensitivity training to reduce infant distress (Milgrom et al., 2010; DeMaster et al., 2019). In preschool-aged children, and specifically those exposed to early adversity (early maltreatment or low SES), electrophysiological measures of cognitive control and selective attention are modified by family-based interventions targeting parent contingency and responsiveness (Bruce et al., 2009; Neville et al., 2013; Isbell et al., 2017). Such “two-generation” intervention programs presumably indirectly enhance children’s neurocognitive development by way of modifying caregiving environments linked to positive developmental outcomes (Fisher et al., 2016).

While there is limited research on the neural effects of modifying children’s language exposure specifically, multiple randomized controlled trials have investigated the efficacy of parent-implemented interventions on children’s linguistic and cognitive development. A meta-analysis of 25 randomized controlled trials targeting children (birth-8 years) with or at risk of language impairment, including 1734 children from lower-SES homes, found that parent-implemented language interventions lead to small-to-modest but reliable improvements in children’s expressive language skills (Heidlage et al., 2020). Additionally, several studies have investigated parent coaching interventions with personalized feedback for families from lower - SES (Leung et al., 2020; Wong et al., 2020), higher-SES (Ferjan Ramirez et al., 2019; Ferjan Ramírez et al., 2020), and mixed-SES (McGillion et al., 2017) backgrounds. Across all of these studies, caregivers who received intervention exposed their children to more words and more conversational turns as measured by child-worn audio-recorders. In studies that additionally examined intervention effects on child language, children whose parents received intervention exhibited more vocalizations (Ferjan Ramirez et al., 2019; Ferjan Ramírez et al., 2020) and larger parent-reported vocabularies (Ferjan Ramírez et al., 2020; McGillion et al., 2017 low-SES families only) and broad language gains (Wong et al., 2020) over the months following intervention. Furthermore, increased conversational turn-taking also correlated with children’s language growth over the following year (Ferjan Ramirez et al., 2019; Ferjan Ramirez et al., 2020; Gilkerson et al., 2017). Although these interventions involved detailed, comprehensive programming, even brief, “light touch” interventions have been effective in increasing the amount of parent-child conversation and measures of children’s expressive language (Leech & Rowe, 2020). The efficacy of input-modifying interventions on children’s language skills raises questions about the underlying neurobiological mechanisms.

The present study aims to fill these gaps by investigating whether and how children’s brain structure and cognition change in response to a family-based intervention designed to enhance the communication environments of children from primarily lower-SES backgrounds. The intervention employed was a direct replication and extension of an interactive, small-group program designed to improve preschool children’s cognition through a variety of parent-child communication practices (Neville et al., 2013). In the original study, families in the intervention group demonstrated more balanced (versus parent-dominated) adult-child verbal turn-taking during a free-play session. Children also demonstrated intervention-related increases in receptive language skills and electrophysiological measures of selective attention (Neville et al., 2013; Isbell et al., 2017), but the mechanisms underlying these cognitive improvements have not been investigated. The present study used naturalistic measures of children’s real-world language exposure to measure changes in children’s language experience, and related this change to growth in children’s verbal and nonverbal cognition and whole-brain cortical thickness. We specifically focused on structural cortical measures to investigate experience-related neuroanatomical plasticity during a developmental period of typical cortical thinning (Remer et al., 2017).

We hypothesized that families who completed the intervention would exhibit increased conversational turn-taking and improvements to children’s language scores. Further, we hypothesized that cortical thickening in left hemisphere perisylvian regions would mediate the relationship between changes in turn-taking and language development. Anticipating individual variability in treatment response (i.e., in how much parents used intervention strategies), we also examined relationships amongst longitudinal changes in language experience, cognitive assessments, and cortical thickness both within and across groups.

## Materials and Methods

### Participants

Participants include 52 children (32 male, 20 female) enrolled in either pre-kindergarten (*n*=14) or kindergarten (*n*=38), aged 4 years, 5 months to 7 years, 1 months (*M*=5.79 years, *SD*=0.60 years). Because recruitment occurred at the family level, the sample comprised 50 families, including one pair of twins and one sibling pair aged 1 year apart. No results of interest change if either child in each sibling pair is excluded. Participants were a subset of participants from a larger intervention study (Romeo & Leonard, et al., under review), which included 107 children (101 families) who were recruited from eight over-subscribed public charter schools in metropolitan Boston that serve primarily lower-income families. All participants in the larger study completed an in-school battery of cognitive assessments (nonverbal IQ, executive functioning, and some language measures). All families from the larger study were then invited to participate in additional in-lab cognitive assessments (more comprehensive language measures), parent surveys, neuroimaging, and home language recording. The present sample includes all families who participated in the additional in-lab assessments. All demographic measures, descriptive statistics, and inferential statistics reported here apply to the sub-sample (n=52) who participated in in-lab assessments. There were no demographic differences between families who participated in the supplemental in-lab sessions and those that did not (all *p* > .29). Two children (both in the control group) did not return for the in-lab post-test session but did complete the in-school post-test session. Figure 1 illustrates the sample and subsample structure. A parent or legal guardian provided written informed consent, all children provided verbal informed assent, and all procedures were approved by the Massachusetts Institute of Technology institutional review board.

**Figure 1.**
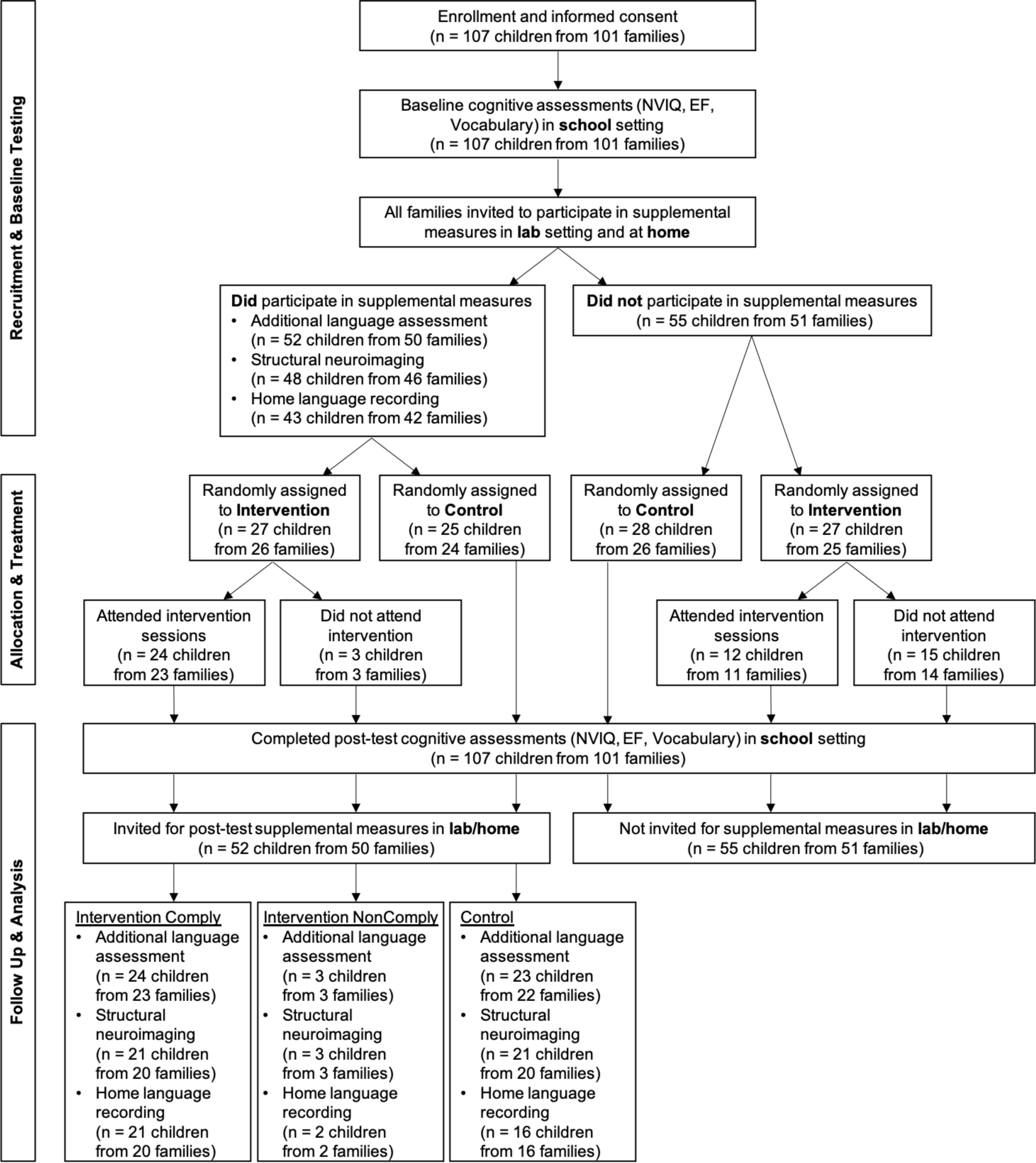
Flowchart illustrating the order and usable sample size of pre- and post-test measures, group assignment, and treatment compliance. NVIQ = Nonverbal IQ and EF = Executive Functioning assessments.

A primary caregiver reported the child’s demographics. Race/ethnicity (“select all that apply”) included *n*=5 White/Caucasian, *n*=25 Black/African American, *n*=29 Hispanic/Latinx, *n*=1 Asian, *n*=1 Native American/American Indian, *n*=2 Native Hawaiian/Pacific Islander, and *n*=1 Other (self-identified as Cape Verdean). Nine participants selected more than one race/ethnicity, which are included in the counts above. Although race/ethnicity are reported for descriptive purposes, neither is used as an independent variable or covariate based on current best practices, in favor of more proximal potential explanatory factors (APA, 2019; Helms et al., 2005). Thirty-four children lived with two parents/caregivers, 17 with one parent/caregiver (of which 7 split time with a second parent), and 1 with a non-parental legal guardian. Parental education (average of both parents, if a two-parent household) varied from 6^th^ grade to advanced degree, and was ordinally coded into five bins: less than high school (*n*=7), high school (*n*=20), some college (*n*=13), four-year college degree (*n*=10), or advanced degree (*n*=2), such that the median was a high school education. In the region where the study was conducted, the median education is a college degree. Annual total household income ranged from less than $6,000 (inclusive of benefits) to $160,000 with a median of $53,500 (*M*=$61,073, *SD*=$42,172), and income-to-needs based on family size and the federal poverty level ranged from 0.24-6.68 (*Med*=2.38). Thirty-eight children (73%) qualified for free or reduced price lunch (household incomes under 185% of the federal poverty level). However, given the high cost of living relative to federal guidelines, participants were additionally classified by regional HUD Income Limits (https://www.huduser.gov/portal/datasets/il.html) into five ordinal bins: “extremely low income” (n=14), “very low income” (n=9), “low income” (*n*=11), “below median income” (n=8), and “above median income” (n=10). The 5-point ordinal scales for parental education and regional income were highly intercorrelated (Spearman’s *ρ*=.638, *p*<.001), but were considered separately in all SES analyses because of potentially differing relationships with child development (Duncan & Magnuson, 2012). All children were native English speakers, though twenty-four (46%) children were reported to know/speak an additional language with moderate proficiency (*n*=16 Spanish, *n*=6 Haitian Creole, *n*=2 Cape Verdean Creole).

In effort to establish a scalable intervention model, all children from the target grades in each school were invited to participate in the intervention. The only inclusion criteria were that parents felt comfortable communicating in either English or Spanish (self-determined). The only exclusion criterion for in-lab participation was MRI contraindications (e.g., non-removable metal). Thus, the present sample of children includes eight children who might otherwise be excluded from language studies, including six children with Individualized Education Plans for speech/language, one child born premature (29 weeks gestation), and one child taking medication for Attention Deficit Hyperactive Disorder (methylphenidate). Children were otherwise typically developing. No reported effects change if developmental typicality is covaried. Additionally, because recruitment occurred at the family level (as opposed to the child level), two families each had two children that participated. One family with twins (1 boy, 1 girl) was in the intervention group and completed all measures, and one family with siblings aged one-year apart (both girls) was in the control group and chose not to complete any home audio

### Study Timeline

Families were recruited over a three-month period preceding the start of the intervention. All children whose parents consented to participate in the larger intervention study (n=107 children from 101 families) completed a pre-test and post-test battery of cognitive assessments in the child’s school during a pullout session during school hours. All participating families were also invited to participate in a supplemental in-lab session which included additional language assessments, parent surveys, neuroimaging, and home language recording, and 52 children from 50 families (49%) did so. Children completed the same battery of assessments before (pretest) and after (posttest) the intervention period. The intervention period lasted 10 weeks (8 core curriculum weeks, 1 introductory week, and 1 spring break week). In-school testing occurred in the two weeks immediately preceding and following the intervention period, and in-lab testing occurred simultaneously in the 6 weeks immediately preceding and following the intervention period (overlapping with in-school testing). Home audio recordings typically occurred the weekend following the in-lab session (*M* = 8.50 days after, *SD* = 4.75 days) and never occurred during the intervention time-period. No time differences between sessions (in-school, in-lab, or home audio) or between pre-post timepoints differed between assigned groups (all *p*>.2). Random assignment occurred after the completion of all pretesting, but because scoring had not been completed, it did not account for any pretest measures. See Figure 1 for a flowchart of the depicting the timeline of data collection, group assignment, and the sample size for each measure.

### Cognitive Assessments

Children completed a battery of standardized behavioral assessments to characterize verbal skills, nonverbal intelligence, and executive functioning. All assessments were administered and scored by researchers blind to participants’ group assignment. All measures except for the *Clinical Evaluation of Language Fundamentals* (CELF-5) (Wiig et al., 2013) were completed in school as a part of the larger intervention study, while the CELF-5 was completed in lab immediately preceding the neuroimaging session (see Figure 1 for timeline). Descriptive statistics of measures are presented in Table 1.

**Table 1.** Baseline scores and intervention effects on language environment and cognition.

| Measure | Pre-Test | | Post-Test | | Time*Group Interaction<br>$\eta^2$ ($p$) |
| --- | --- | --- | --- | --- | --- |
|  | Intervention<br>M (SD) [n] | Control<br>M (SD) [n] | Intervention<br>M (SD) [n] | Control<br>M (SD) [n] |  |
| Adult Words per Hour | 1106.50<br>(381.61) [23] | 1258.81<br>(489.72) [18] | 1325.27<br>(419.16) [21] | 1183.68<br>(451.35) [16] | .076 (.098)<sup>†</sup> |
| Conversational Turns per Hour | 47.03<br>(18.88) [23] | 55.77<br>(21.79) [18] | 51.63<br>(20.02) [21] | 48.71<br>(20.73) [16] | .159 (.014)* |
| Child Vocalizations per Hour | 206.71<br>(83.40) [23] | 222.84<br>(91.39) [18] | 216.63<br>(85.71) [21] | 233.59<br>(111.72) [16] | .002 (n.s.) |
| Language Composite | 98.40 (16.43)<br>[24] | 100.18 (15.73)<br>[25] | 101.83 (14.54)<br>[24] | 103.88 (13.36)<br>[25] | .000 (n.s.) |
| Nonverbal IQ Composite | 9.42 (2.23)<br>[24] | 9.93 (2.23)<br>[25] | 10.60 (2.18)<br>[24] | 10.45 (2.26)<br>[25] | .066 (.074)<sup>†</sup> |
| Executive Functioning Composite | -.15 (.63)<br>[24] | 0.05 (.77)<br>[25] | 0.05 (.54)<br>[24] | -.09 (.86)<br>[25] | .096 (.031)* |

<sup>†</sup> $p < 0.1$ , \* $p < 0.05$

Verbal assessments included the *Peabody Picture Vocabulary Test* (PPVT-4) (Dunn & Dunn, 2007), which measures receptive vocabulary knowledge, and four subtests comprising the Core Language Score (CLS) of the CELF-5, which includes Sentence Comprehension (measures general receptive language), Word Structure (measures expressive morphosyntax), Formulated Sentences (measures expressive semantics and morphosyntax), and Recalling Sentences (measures verbal memory and general expressive language). Because the CELF-5 is age-normed for children aged 5-21 years, standard scores are unavailable for n=7 participants who were 4 years-old at the time of assessment. This assessment was chosen because pilot testing with language assessments designed for younger children revealed ceiling effects for older students and presumably would have allowed for minimal growth after intervention. PPVT-4 data were invalid for one child at pretest and another child at posttest; all CELF-5 data were valid. Standard scores from the two language assessments were averaged to create a composite language score, except where only one standard score was available.

Nonverbal cognitive assessments included three subtests of the *Wechsler Preschool and Primary Scale of Intelligence* (WPPS-IV) (Wechsler, 2012), which included Matrix Reasoning (measures nonverbal fluid reasoning), Picture Memory (measures nonverbal working memory), and Bug Search (measures nonverbal processing speed). Age-normed scaled scores from the three WPPSI-IV subtests were averaged to create a composite nonverbal IQ score. Bug Search data were invalid for one child at pre-test and another child at post-test, so for these two children, the other two measures (Matrix Reasoning and Picture Memory) were averaged to create a nonverbal IQ composite score.

Executive functioning was measured by the Head-Knees-Toes-Shoulders (HKTS) task (McClelland et al., 2014) and the Hearts and Flowers (HF) version of the dots task (Davidson et al., 2006), both of which require children to follow directions that involve changing instructions to tap inhibition and cognitive flexibility skills. In HKTS, children are first asked to touch their head when told to touch their toes and vice versa (10 trials), then two more body parts (knees and shoulders) are added to the command sequence and opposite commands continue (10 trials), and finally, the rules are changed (i.e., head goes with knees, and toes go with shoulders, 10 trials). Children received six practice items before each block. A score of 0 is given for an incorrect response, 1 for a self-correct, and 2 for a fully correct response, bringing the maximum score to 60. HF was presented on a touch-screen tablet via the Presentation program by Neurobehavioral Systems (Berkeley, CA). In each trial, a red heart or flower would appear on the right or left side of the screen. First, children are told to press the button on the same side as the stimulus (congruent block, 12 trials), then to press the button on the opposite side of the stimulus (incongruent block, 12 trials), and finally to press the same side as heart but the opposite side of a flower (mixed block, 49 trials). For all conditions, stimuli were displayed for 500 msec with 1500 msec to respond and an interstimulus interval of 500 msec. Children received up to 12 practice trials before each block. The two outcome measures include accuracy (percent correct) and reaction time on correct trials (in milliseconds) across the incongruent and mixed blocks combined. Any response faster than 200 msec was considered to be anticipatory (Davidson et al., 2006) and thus were excluded from analyses, and the first trial of the mixed block was also excluded. HKTS data was invalid for two children at posttest; HF data was valid for all children. HKTS total score, HF accuracy, and HF reaction time (reversed) were z-scored and averaged to form an executive functioning composite score.

### Parent Survey Measures

At both pre and post in-lab sessions, parents completed questionnaires assessing their caregiving experience, their child’s behavior, and the parent-child and family relationships. One family did not complete the survey measures at either time point. Two measures were analyzed in the current study. The Perceived Stress Scale (PSS), 10 Item version (Cohen et al., 1983; Cohen & Williamson, 1988) measures the degree to which parents appraise their lives to be stressful and taps how unpredictable, uncontrollable, and overloaded parents feel. The Parenting Relationship Questionnaire (PRQ) is a component of the *Behavior Assessment System for Children, Third Edition* (BASC-3; Reynolds & Kamphaus, 2015) which assesses the parent’s perspective on the parent-child relationship. Two subscales were analyzed: the Parenting Confidence (PC), which measures parents’ comfort, control, and confidence when making parenting decisions, and Discipline Practices (DP), which measures consistent discipline alongside a belief that rule adherence is desirable. The PRQ has different versions for parents of children 5 years and younger (PRQ Preschool version) and 6 years and older (PRQ Child/Adolescent version). The PC subscale has 7 items in the preschool version, and the same items plus one additional item in the child version. The DP subscale has the same 9 items in each version. Parents were administered the relevant version for their child’s age on the day of assessment, and norm-referenced standard scores were analyzed.

### Neuroimaging Data Acquisition and Analysis

Child participants completed identical neuroimaging sessions at pre-test and post-test. First, children were acclimated to the MRI environment and practiced lying still in a mock MRI scanner. Data were then acquired on a 3 Tesla Siemens MAGNETOM Trio Tim scanner equipped for echo planar imaging (EPI; Siemens, Erlangen, Germany) with a 32-channel phased array head coil. A whole-head, high-resolution T1-weighted multi-echo MPRAGE (van der Kouwe et al., 2008) structural image was acquired while children watched a movie of their choice, using a protocol optimized for movement-prone pediatric populations (Tisdall et al., 2012) (TR = 2530 ms, TE = 1.64 ms/3.5 ms/5.36 ms/7.22 ms, TI = 1400 ms, flip angle = 7°, resolution = 1-mm isotropic). The scan session also included a story-listening task, resting state, and a diffusion weighted scan (all collected after the structural scan), but structural measures were chosen as current measure of interest because it had the largest number of participants with two timepoints of usable data. All neuroimaging took place at the Athinoula A. Martinos Imaging Center at the Massachusetts Institute of Technology, and was conducted by researchers blind to participants’ assigned group.

T1-weighted images from both time points were visually inspected for image quality (based on a guide of artifacts associated with motion) by two trained observers blind to all participant measures. Additionally, image quality was objectively quantified by QoalaT (Klapwijk et al., 2019), a supervised learning tool for evaluating structural MRI images. One participant’s imaging data from timepoint 1 was dropped for failing both subjective and objective quality control. Additionally, three participants chose not to complete imaging at both timepoints, one participant chose not to complete imaging at the second timepoint, and two participants did not return for the second in-lab session which included imaging (timepoint 1 n=48, timepoint 2 n=46, longitudinal n=45). QoalaT scores of included participants were highly correlated between timepoints (*r*(43)=.697, *p*<.001) indicating consistent within-participant scan quality. Quality ratings were not significantly correlated with any cognitive or demographic variables, and did not differ by intervention group (all *p>*.1).

Surface-based cortical reconstruction was conducted with FreeSurfer v5.3.0 (Fischl 2012). Two trained researchers manually edited pial and white matter surfaces as needed, and a third researcher confirmed the accuracy of the final surfaces. T1 images from both timepoints were processed with FreeSurfer’s longitudinal stream (Reuter et al. 2012). This process estimates average participant anatomy by creating an unbiased within-participant template space (Reuter and Fischl 2011) using a robust, inverse consistent registration (Reuter et al. 2010) and resampling cross-sectional images to the template. Within-participant templates and resampled cross-sectional reconstructions were manually edited and checked (as above), and then warped to a standard brain (fsaverage) and smoothed with a 15-mm full-width half-maximum kernel. General linear models were constructed with a dependent variable of symmetrized percent change (SPC), which is the rate of change at each surface vertex with respect to the average thickness across both timepoints. SPC is more robust than rate of change or simple percent change, which refer to change only in terms of the first measurement. All analyses controlled for participant age and gender. Whole-brain analyses were cluster-corrected for multiple comparisons using a Monte Carlo simulation with 10,000 repetitions with absolute cluster-forming *p*<.01 and cluster-wise *p*<.001, Bonferroni adjusted for both hemispheres (Hagler et al., 2006).

### Home Audio Recordings

At each in-lab session, parents were introduced to the Language Environment Analysis (LENA) recording system and how to use it (Gilkerson et al., 2017). Following both pretest and posttest in-lab sessions, parents were mailed two LENA Pro digital language processors (DLPs), which are small 2-ounce digital recorders that fit in a shirt pocket and store up to 16 hours of digitally recorded audio (Ford et al., 2008). Parents were instructed to collect two full-day recordings from consecutive days when the child was not in school (typically the Saturday and Sunday following the in-lab session), beginning when the child wakes up, and to conduct their days as normal. Due to technical difficulties, four total recording timepoints contained only one day of recording. After families mailed back the DLPs, LENA software employed validated speech-identification algorithms (Xu et al., 2009) to automatically segment the audio and provide estimates of 1) the number of adult words spoken to the child (AWC), 2) the number of child vocalizations at the utterance level (CVC), and 3) the number of adult-child conversational turns (CTC), defined as a discrete pair of an adult vocalization followed by a child vocalization, or vice versa, with no more than 5 seconds pause between the two. Parents of 9 children (including a sibling pair) either declined to participate in the LENA recording or provided unusable recordings at the first time point (valid n=43), plus an additional 4 children at the posttest timepoint (valid longitudinal n=39: 21 intervention, 16 control, 2 non-comply).

Although parents were instructed to keep the recorder on and with the child for full days, the recorder was occasionally turned off (sometimes by accident) or removed from the child (e.g., during bath time, sports practices, etc.). A combination of parent-reported activity logs and the raw audio recordings were screened for times when the recorder was off, not on the child’s person, or the child was asleep, and these segments were removed from analysis. Audio file screening was completed by a researcher blind to all participant demographics, assessment scores, and group assignment. Mean hourly AWC, CVC, and CTC were calculated by dividing each total count metric by the total duration of usable recording hours across both days (mean analyzed hours per timepoint = 21.41 hours, SD = 6.10 hours).

### Intervention

Participants were randomly assigned at the family level to either attend a family-based intervention program or a no-contact, “business-as-usual” control group. Random assignment was done within the full in-school sample (n=107, see Participants section above), stratifying for school and whether participants had completed the additional in-lab session. Within the present sample, n=27 participants (including one sibling pair) were assigned to the intervention, and n=25 participants (including one sibling pair) were assigned to the control group. Intervention and control groups did not differ on any demographic measure, or baseline environmental or cognitive measures (all *p*>.19). Three participants (no siblings) assigned to the intervention group did not attend the intervention (“non-compliers”) and are excluded from analyses investigating intervention efficacy, but because they completed all measures at both timepoints, they are included in longitudinal correlational analyses. Intent-to-treat analyses including these children in the intervention group are reported in the Supplement.

The intervention was a near-exact replication of the Parents and Children Making Connections – Highlighting Attention (PCMC-A) program designed by Neville and colleagues (Neville et al., 2013), and this specific replication is described in detail elsewhere (Romeo & Leonard, under review). Briefly, PCMC-A consists of nine (1 introduction, 8 content) weekly small group sessions (4-5 families per group). Trained Parent Facilitators with extensive family engagement experience led parents/caregivers in a two-hour interactive curriculum of strategies to target family stress regulation, contingency-based discipline, parental responsiveness and language use, and facilitation of child attention. Simultaneously, trained Child Facilitators with extensive child education experience led children in a parallel 45-minute child curriculum involving multi-sensory activities and games aimed to increase self-regulation of attention and emotion states. At the conclusion of the child curriculum, children received free childcare until the conclusion of the parent session. Between sessions, parents were asked to try the strategies at home, and they received weekly 15-minute check-in phone calls from group facilitators to discuss implementation and effectiveness of the strategies. Parents assigned to the intervention had the choice of attending groups led in either English (n=20) or Spanish (n=4), while the child component of the intervention was delivered in English, to match the children’s language of instruction at school. Make-up sessions were available if a family missed a scheduled session. During sessions, families received a free meal, child-care for any siblings or other children under the adults’ care, a travel reimbursement, and the opportunity to win additional incentives. Facilitators completed a fidelity protocol designed by PCMC-A staff to ensure reliability of the curriculum. Parent and child facilitators held weekly meetings with the intervention designers to debrief on previous sessions, ensure faithful and consistent curriculum delivery, and prepare for upcoming sessions. The discussion protocol included confirming principal intervention features, including both curriculum content (e.g., correct information, time allotted for each section) and delivery process (e.g., the facilitator’s affect, promotion of discussion, and handling of challenges). No curriculum deviations were noted in any of the weekly meetings.

While the PCMC-A curriculum includes eight distinct topics primarily designed to improve children’s self-regulation skills, many of the lessons explicitly target parent-child communication practices and language use. For example, one module focused on optimizing learning through play by instructing parents to follow the child’s interests/attention and to balance conversation during play by letting child start talking first, taking a turn after the child (using social language to model new vocabulary and concepts), and counting to five if they realize they are doing all the talking without letting the child engage. Another module instructs parents to use “meaningFULL” language which incorporates open-ended questions that encourage a child response (as opposed to meaningless questions to which a child’s response is irrelevant), and to increase specific verbal praise in response to children’s actions. Additional modules target using clear communication that is pleasant and respectful, is age-appropriate in content and structure, makes use of routines, maintains the child’s attention, and verbalizes thinking, emotions, and actions. Throughout all lessons, parents are encouraged to be more aware of their language use, tune in to their child’s verbal and nonverbal communication, and aim to balance communication in parent-child connections. At the start of each session, facilitators reviewed the previous week’s strategies and engaged parents in discussion of their experiences to ensure uptake.

### Analysis Plan

First, we evaluated the effects of the intervention on home language, child cognitive measures, and brain plasticity through repeated measures general linear models with time as the within-subjects variable and group as the between-subjects variable. Effects of the intervention were determined by significant time by group interactions. For significant effects, we also report effect sizes (partial eta squared) and robust *p-*values calculated with 10% trimming. Second, we investigated how longitudinal changes in the language environment related to changes in children’s cognitive scores and brain structure by examining correlations of pre-to-post changes within the intervention group (to examine variability in treatment response), and in both groups combined (to examine longitudinal changes irrespective of treatment condition). For significant effects, we also report robust *p-*values calculated with 10% trimming. This analysis has the advantage of maximizing statistical power at the expense of a causal design. Third, we estimated bootstrapped mediation models (10,000 iterations) with 95% bias-corrected confidence intervals to investigate mechanistic relationships between intervention group, changes in turn-taking, changes in cognitive scores, and neuroplasticity. Finally, we explored whether alternative explanatory measures, namely parents’ perceived stress and parenting confidence, were affected by the intervention and explained cognitive effects better than turn-taking changes, using the above methods plus robust multiple regression with Huber weighting.

## Results

At baseline, the two groups did not significantly differ on any demographic, cognitive, experiential, or neural measures (all *p*s > .19). Baseline correlations between SES, language environment, and child cognitive scores (collapsed across groups) are reported in Supplementary Table 1.

### Intervention Effects on Experiential and Cognitive Measures

A significant time by group interaction (Figure 2, Table 1) revealed that families who attended the intervention exhibited a greater increase in average hourly conversational turns than families in the control group [*F*(1,35)=6.63, *p*=.01, trimmed *p*=.04, partial *η*^2^=.16; mean change intervention = +4.24, control = −7.01], in addition to a marginal increase in the average hourly adult words [*F*(1,35)=2.89, *p*=.098, trimmed *p*=.11, partial *η*^2^=.08; mean change intervention = +189.83, control = −55.25]. The effect of the intervention on conversational turns remained significant when controlling for sex, age, SES, language of intervention delivery, or child bilingualism (all *p*<.05), and no demographic measure moderated intervention effectiveness. There were no group differences in changes in the number of child vocalizations (*p*=.78).

**Figure 2.**
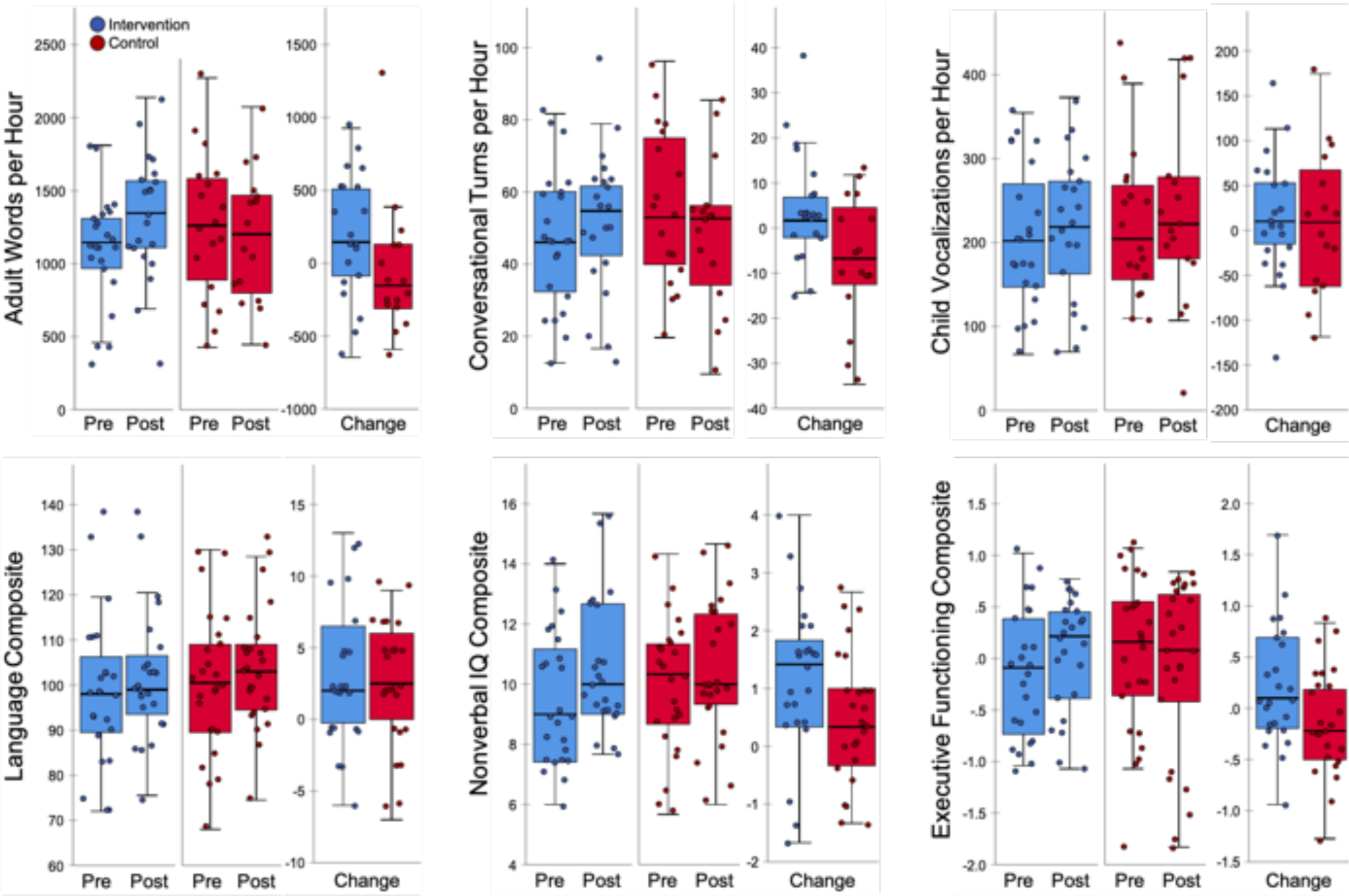
Pre-to-post changes in language environment measures (top) and standardized cognitive assessments (bottom), split by intervention group (intervention = blue, control = red). Change scores represent posttest minus pretest scores. Each dot represents a participant, overlaid on boxplots representing subgroup median (dark center line), first and third quartiles (box edges), and the maximum and minimum values no larger than 1.5 times the interquartile range (whiskers). See Table 1 for corresponding descriptive and inferential statistics.

A main effect of time on the language and nonverbal composites revealed that all children on average exhibited increases in language scores [*F*(1,47)=11.88, *p*=.001, robust *p*=.002, partial *η*^2^=.20; mean change = +3.49] and nonverbal-IQ scores [*F*(1,47)=22.41, *p*<.0001, robust *p*=.0004, partial *η*^2^=.32; mean change = +.79]. There was no main effect of time on executive functioning. However, a significant time by group interaction (Figure 2, Table 1) revealed that children whose families attended the intervention exhibited a relative increase in executive functioning scores compared to peers in the control group [*F*(1,47)=4.97, *p*=.03, robust *p*=.03, partial *η*^2^=.10; mean change intervention = +.21; control = −.14], in addition to a marginally greater increase in nonverbal-IQ scores [*F*(1,47)=3.34, *p*=.07, robust *p*=.19, partial *η*^2^=.07; mean change intervention = +1.17 control = +.52] (however, neither of these effects were significant in the full-intervention study sample, Romeo & Leonard et al., under review). There was no time by group interaction on language scores, as children in both groups increased comparably (mean change intervention = +3.72; control = +3.48). Because of these limited group-level effects on cognition, the remainder of results focus on individual differences in changes over time.

### Longitudinal Changes in Language and Cognition

To determine how longitudinal changes in language environment related to changes in children’s cognitive scores, we examined pre-to-post changes both within the intervention and control groups, and in both groups combined. Collapsing across groups, the magnitude of change in conversational turns was significantly correlated with the magnitude of change in language scores [*r*(37)=.37, *p*=.02, robust *p*=.01], nonverbal-IQ scores [*r*(37)=.39, *p*=.01, robust *p*=.01], and executive-functioning scores [*r*(37)=.48, *p*=.001, robust *p*=.002] (Figure 3, Table 2). There were no significant relationships between changes in adult words or child vocalizations with any cognitive measure (all *p*>.1). Additionally, group significantly moderated the relationship between changes in conversational turns and language scores [*t*(35)=2.34, *p*=.03, robust *p*=.03], such that children in the intervention group exhibited a strong positive correlation between change in conversational turns and change in language scores [*r*(19)=.57, *p*=.007, robust *p*=.005], while children in control group did not show any relationship. There was no moderation by group for nonverbal-IQ or executive functioning scores. Notably, no demographic measure (sex, age, parental education, family income, bilingualism, or developmental typicality) was associated with changes in any cognitive or LENA measure, either within groups or across the entire group.

**Figure 3.**
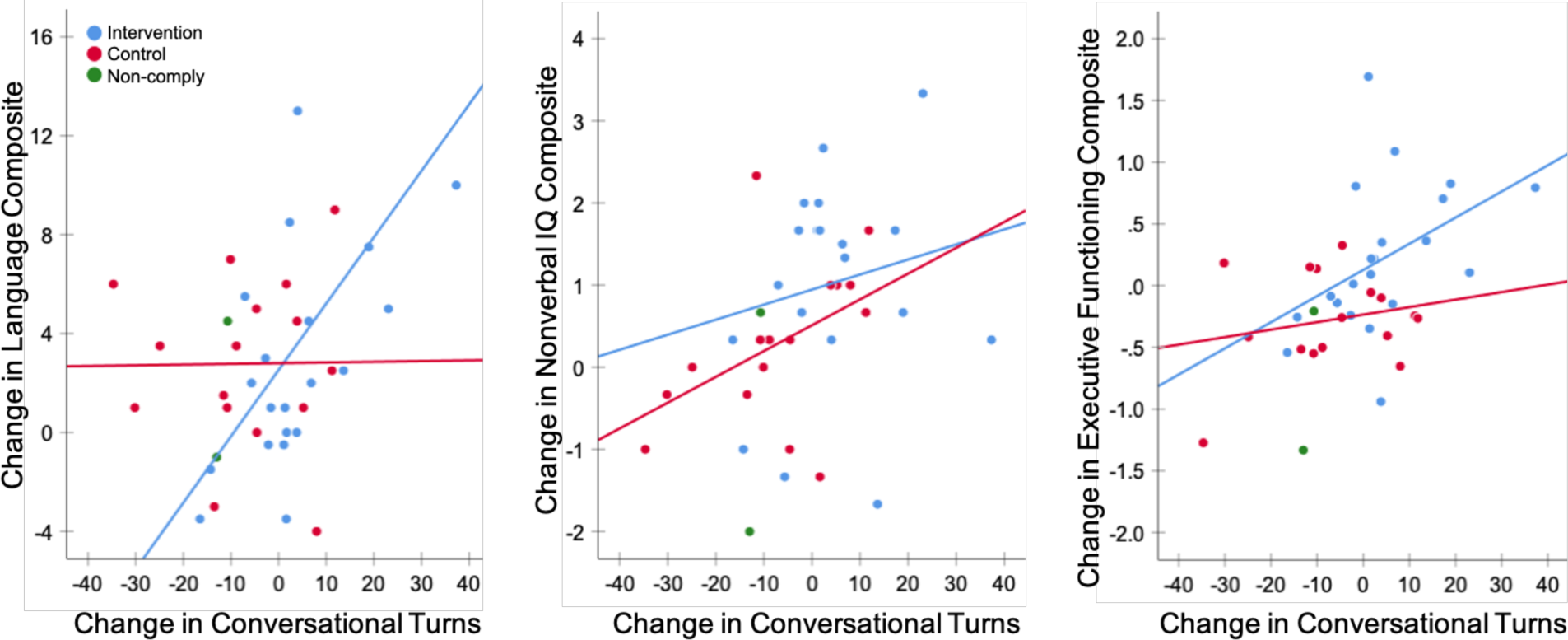
Scatterplots of the pre-to-post change in standardized cognitive measures as a function of change in conversational turns. Colored lines represent the least-squares regression trendline in the intervention (*n*=21, blue) and control (*n*=16, red) groups, excluding two participants assigned to the intervention who did not attend (green). Full sample correlations were significant for all three cognitive measures (see Table 2), and the interaction between groups was significant for language scores (see text).

**Table 2.** Correlations amongst language environment and cognition change scores.

|  | AW | CT | CV | Lang | NVIQ | EF |
| --- | --- | --- | --- | --- | --- | --- |
| Change in Adult Words (AW) | 1 |  |  |  |  |  |
| Change in Conversational Turns (CT) | .44** | 1 |  |  |  |  |
| Change in Child Vocalizations (CV) | 0.06 | .65*** | 1 |  |  |  |
| Change in Language Scores (Lang) | 0.21 | .37* | 0.07 | 1 |  |  |
| Change in Nonverbal IQ Scores (NVIQ) | 0.11 | .39* | 0.21 | 0.18 | 1 |  |
| Change in Executive Functioning Scores (EF) | 0.28<sup>†</sup> | .48** | 0.18 | 0.05 | 0.18 | 1 |

<sup>†</sup> $p < 0.1$ , \* $p < 0.05$ , \*\* $p < 0.01$ , \*\*\* $p < 0.001$
Correlation matrix of change scores in LENA language environment measures and standardized cognitive assessments, collapsed across both intervention (including comply and non-comply) and control groups.
Change scores represent posttest minus pretest scores. Valid pre-post LENA $n=39$ and cognitive $n=52$ .

Next, mediation models were estimated to determine whether the change in conversational turns mediated the intervention effects on cognition. Modern mediation approaches recommend estimating indirect effects even in absence of direct effects (Hayes, 2009; Rucker et al., 2011). The change in conversational turns significantly mediated relationships between assigned group and changes in language scores [indirect effect = 1.55 (.08, 3.64)] and executive-functioning scores [indirect effect = 0.15 (.01, .37)], but not nonverbal-IQ scores [indirect effect = .28 (−.04, .67)].

### Intervention Effects on Neuroplasticity

There were no significant between-group differences in cortical plasticity from first to second timepoint at the whole brain level. However, across all participants combined, the magnitude of change in conversational turns was significantly, positively correlated with cortical plasticity in two left hemisphere perisylvian clusters (Figure 4, Table 3). Specifically, increases in conversational turns were associated with cortical thickening in a cluster in the left inferior frontal gyrus (IFG), extending over pars opercularis and pars triangularis, as well as a cluster in the left supramarginal gyrus (SMG) at the intersection of the inferior parietal lobe and the posterior temporal lobe. To aid in interpretation of these correlations, participants were divided into two groups by a median split on the magnitude of change in conversational turns, which also corresponded to a division into those with increased turns and those with decreased turns. Children who experienced increased turns exhibited a mean *thickening* of 0.29% (8.41 μm) in the IFG cluster and of 0.24% (6.96 μm) in the SMG cluster, while children who experienced a decrease in turns exhibited a mean *thinning of* 0.36% (10.44 μm) in the IFG cluster and pf 0.24% (9.86 μm) in the SMG cluster. There were no significantly correlated regions in the right hemisphere, and there were no regions where adult words or child vocalizations were significantly correlated with plasticity.

**Figure 4.**
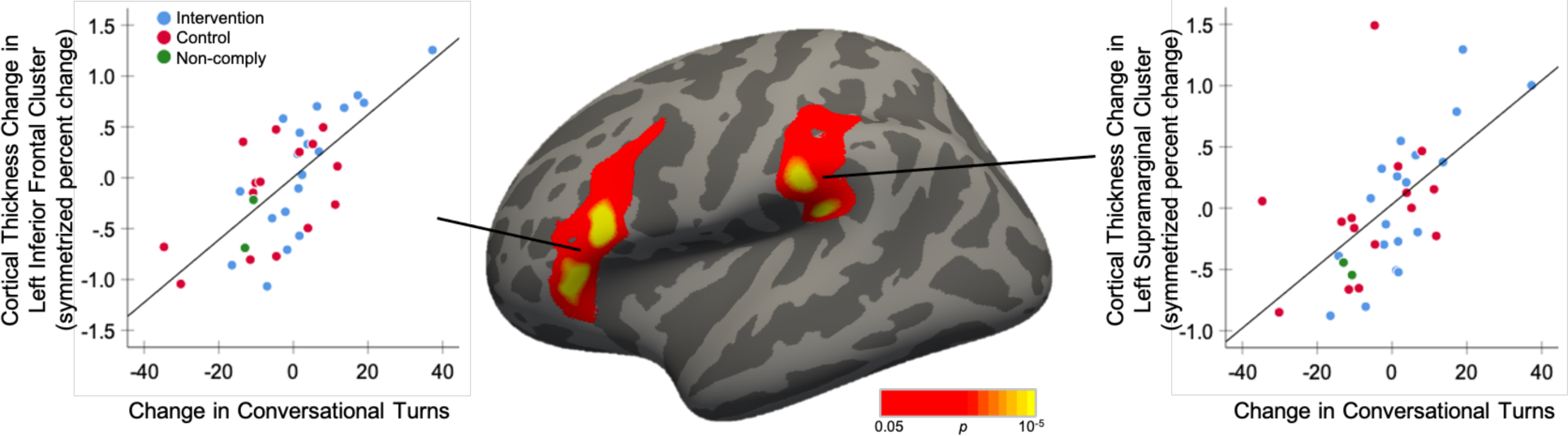
Clusters where changes in cortical thickness are significantly correlated with changes in children’s experienced conversational turns (*n*=36 children, collapsed across groups, controlling for age and gender). Scatterplots represent the average symmetrized percent change in cortical thickness across each vertex per cluster as a function of the pre-to-post changes in conversational turns. Trendlines are for illustrative purposes only.

**Table 3.** Cortical thickness changes associated with changes in conversational turns.

| Region of Cluster<br>(Desikan-Killiany<br>Atlas) | Approximate<br>Brodmann<br>Areas | Area of<br>cluster<br>(mm<sup>2</sup>) | Peak<br>Significance<br>($-\log_{10} p$) | Peak MNI Coordinate | | | Cluster-wise $p$ |
| --- | --- | --- | --- | --- | --- | --- | --- |
|  |  |  |  | x | y | z |  |
| Left inferior frontal<br>gyrus (IFG) | 44, 45 | 1912.32 | 5.450 | -50.8 | 21.3 | 18.0 | 0.00020 |
| Left supramarginal<br>gyrus (SMG) | 40 | 1579.73 | 4.578 | -42.3 | -35.7 | 21.9 | 0.00020 |

Summary of clusters in which symmetrized percent change in cortical thickness are significantly correlated with changes in children's experienced conversational turns ( $n=36$ children, collapsed across groups, controlling for age and gender). Regional labels are derived from the Desikan-Killiany gyral-based atlas (Desikan et al., 2006).

Mediation models were estimated to assess mechanistic relationships among the intervention, changes in turn-taking and cognition, and neuroplasticity. First, changes in conversational turn taking significantly mediated the relationship between the intervention and plasticity in both the IFG [indirect effect = 0.28 (.02, .60)] and SMG [indirect effect = 0.24 (.01, .53)] clusters. Additionally, plasticity in the SMG cluster mediated the relationship between conversational turn-taking changes and language score changes [indirect effect = .08 (.01, .18), Figure 5], with SMG plasticity accounting for 57% of the total relationship between conversational turn-taking increases and language growth. The IFG cluster did not significantly mediate the turns-language relationship, and neither cluster mediated relationships with nonverbal IQ or executive functioning.

**Figure 5.**
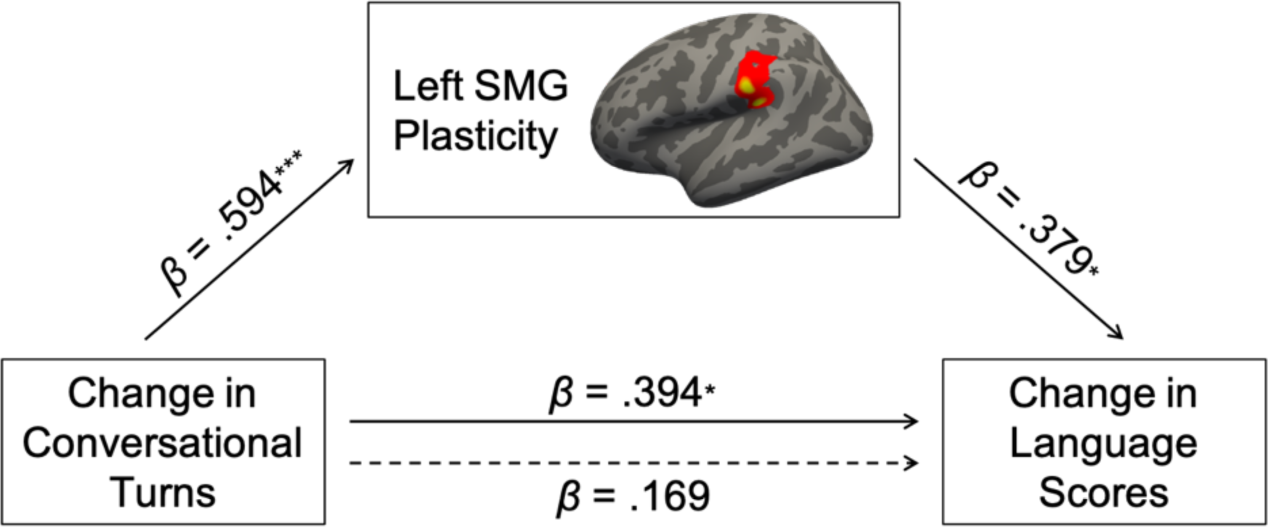
Mediation model showing the relationship between changes in conversational turns and language scores, as mediated by the average cortical thickness change in the cluster in the supramarginal gyrus (SMG). The dotted arrow represents the direct effect after accounting for SMG plasticity. *β* coefficients represent standardized regression coefficients, \**p <* 0.05, \*\**p <* 0.01, \*\*\**p <* 0.001.

### Alternative Explanatory Measures

Because the multifaceted intervention targeted multiple family processes beyond language use, we also examined whether changes in parental stress, parenting confidence, or discipline practices could explain intervention effects in addition to or instead of changes in language experience. Significant time by group interactions, indicating intervention effects, were seen for both perceived stress [*F*(1,44)=5.36, *p*=.025, trimmed *p*=.009, partial *η*^2^=.11] and discipline practices [*F*(1,44)=6.23, *p*=.016, trimmed *p*=.02, partial *η*^2^=.12], but not for parenting confidence. Parents who attended the intervention exhibited a significant decrease in perceived stress (M=−3.75 points) and a non-significant increase in reported discipline practices (M=+1.58 points), while parents in the control group exhibited a non-significant change in stress (M=+0.64 points) and a significant reduction in discipline practices (M=-4.68 points). Collapsing across groups, improvements in EF were significantly correlated with reductions in parental stress [*r*(49)=-.42, *p*=.003, robust *p*=.002] and increases parenting confidence [*r*(49)=.42, *p*=.003, robust *p*=.02], and marginally correlated with increases in discipline practices [*r*(49)=.25, *p*=.08, robust *p*=.47]. No parent measures were correlated with changes in children’s language or nonverbal IQ (all *p*>0.7). While changes in parental stress and parenting confidence were correlated with each other [*r*(49)=-.47, *p*=.001, robust *p*=.001], these were not correlated with changes in conversational turn-taking (both *p*>0.6), suggesting independent pathways. A multiple regression model with EF changes as the dependent variable revealed significant unique effects of changes in conversational turn-taking (*β*=.44, *p*=.001, robust *p*=.0003), parental stress (*β*=−.29, *p*=.04, robust *p*=.03), and parenting confidence (*β*=.32, *p*=.02, robust *p*=.03). Finally, the relationship between intervention group and changes in EF was mediated by changes in conversational turns [indirect effect = 0.15 (0.01, 0.37)], and separately by perceived stress [indirect effect = 0.11 (0.01, 0.25)], but not by parenting confidence [indirect effect = 0.02 (−0.12, 0.13)], When investigated simultaneously, only conversational turn-taking significantly mediated the effect of the intervention on EF [turn-taking: indirect effect = 0.28 (0.03, 0.63); perceived stress: indirect effect = 0.17 (−.05, 0.44)].

## Discussion

This study investigated cortical neuroplasticity associated with changes to children’s language environments following a randomized controlled trial of a family-based intervention for children from primarily lower-SES backgrounds. Families who completed the intervention exhibited greater increases in the quantity of adult-child conversational turns compared to a passive control group, regardless of family SES. Although there were limited effects of group on child-level cognitive or neural measures, across both groups combined, the magnitude of conversational-turn change was positively correlated with changes in children’s language scores, nonverbal cognition, and executive functioning. Further, there were significant indirect effects of the intervention on language and EF through conversational turns. Increases in conversational turn-taking also correlated with greater cortical thickening in left inferior frontal and supramarginal cortices, and plasticity in the supramarginal cluster mediated the relationship between conversational turn and language score changes. These results represent the first longitudinal investigation into the mechanisms of neuroplasticity that underlie changes to children’s language environments, and suggest that conversational turns support children’s language development through growth in specific left perisylvian brain regions.

Although the employed intervention covered a wide range of family-based strategies for supporting children’s broader cognitive development, the crux of the curriculum focused on adult-child communication practices, and specifically parental responsiveness and “meaningFULL” language use. The original evaluation of this intervention found that families who completed the intervention demonstrated more balanced (versus parent-dominated) adult-child verbal turn-taking during an 8-minute in-lab play session (Neville et al., 2013). The present study found commensurate results using a different methodology, specifically two days of “real-world” audio recordings of children’s language environments, suggesting that the intervention’s effect on children’s communication experiences extend to more naturalistic circumstances. Importantly, this result remained significant after controlling for family SES (both parental education and household income), suggesting that the intervention is equally effective on increasing conversational turn-taking over the sampled SES range.

Despite the positive effects on children’s language environments, there were limited between group differences in children’s cognitive scores, in contrast with the findings of Neville and colleagues (Neville et al., 2013). Potential reasons for this non-replication include participants being on average one year older; the use of different standardized assessments, including more comprehensive language measures; and differences in the racial/ethnic, geographical, and cultural background of participating families, including more participants from racial and ethnic minority backgrounds. Future research is needed to determine which populations receive the most benefit from this intervention and whether tailoring to specific recipients may yield more widespread positive results in children’s cognition.

Given the goal of this study to highlight mechanisms of experience-driven language development, the most intriguing findings are the relationships between changes in conversational turn-taking and plasticity in children’s cognition and brain structure. Previous studies investigating neural mechanisms underlying language input-output relationships have all been cross-sectional (Merz et al., 2020; Romeo, Leonard, et al., 2018; Romeo, Segaran, et al., 2018), which limits the ability to make causal interpretations. The study extends these results, by revealing indirect effects of an intervention through changes in turn-taking, and by demonstrating associations between *longitudinal* changes in children’s language environments, language development, and structural neuroplasticity. Together, this provides the strongest evidence to date that children’s language experience—and specifically adult-child conversational turn-taking—promotes cortical growth in left perisylvian brain regions supporting children’s language development in early childhood.

The two brain regions that exhibited plasticity associated with conversational turns are well-established as core hubs of the language network. The left IFG, often referred to as Broca’s area, is causally involved in speech production and frequently implicated in receptive language and reading tasks (for review, see Friederici, 2011). A prominent theory suggests that Broca’s area operates as a convergence zone in which the phonological, syntactic, and semantic components of language are unified into a coherent linguistic representation (Hagoort, 2014). Other theories posit that Broca’s area serves a more domain-general function of processing hierarchical structures, of which language is one instantiation (Tettamanti & Weniger, 2006), or that subregions of Broca’s area selectively support language processing and diverse set of attentionally demanding perceptual and cognitive tasks (Fedorenko & Blank, 2020). Indeed, the left IFG is a part of the dorsolateral prefrontal cortex which supports higher-level nonverbal skills such as fluid reasoning (Ferrer et al., 2009), as well as a broader prefrontal network supporting goal-directed behavior and executive functioning skills such as working memory, cognitive control, cognitive flexibility, inhibition, and selective attention (Diamond, 2013; Miller & Cohen, 2001). Conversational turn-taking may engage many of these executive processes, such as utilizing working memory to remember previous statements in the conversation, or inhibiting speech until the previous speaker is done. The finding that changes in conversational turn-taking correlated not only with growth in language growth, but also nonverbal cognition and executive function, suggests that conversational turn-taking may directly support the development of executive processes through either shared or adjacent prefrontal mechanisms. Furthermore, the left SMG, located at the intersection of the inferior parietal lobe and posterior temporal lobe, also subserves aspects of language and nonverbal cognition. The left SMG is frequently considered to be a part of Wernicke’s area (Tremblay & Dick, 2016), which is critically involved in language comprehension, especially as it relates to processing the phonological components of words and recoding phonology to lexical cues (Oberhuber et al., 2016). Furthermore, the left temporoparietal junction, located at the posterior edge of the SMG, is recognized as a core region underlying social cognition, including the ability to reason about others’ thoughts and beliefs, known as theory of mind (Adolphs, 2009), which undergoes significant development during the age range studied (Richardson et al., 2018). Conversational turn-taking necessarily requries social reciprocity and may promote theory of mind processes such as perspective taking when interacting with an interlocutor. In return, some theories posit that language acquisition and growth *requires* social interaction (e.g., Kuhl, 2007; Tamis-LeMonda et al., 2014). The finding that SMG plasticity mediates the relationship between conversational turns and language score changes suggests that social interactive processes may drive language development during this important developmental period. The present study did not directly probe the effects of the intervention on children’s social cognition, so future research is needed to comprehensively explore the complex relationships amongst these cognitive domains during their co-development in preschool period.

The present results add to the growing number of findings that conversational turns are one of the strongest predictors of children’s linguistic, cognitive, and neural development (Ferjan Ramírez et al., 2020; Gilkerson et al., 2018; Hirsh-Pasek et al., 2015; Leech & Rowe, 2020; Merz et al., 2020; Romeo, Leonard, et al., 2018; Romeo, Segaran, et al., 2018; Zimmerman et al., 2009). This raises empirical questions about what component of conversational turn-taking confers such benefits to child development, over and above other measures of language input. Children’s language exposure is multifaced and includes linguistic (e.g., syntax), social-interactive (e.g., contingency), and conceptual (i.e., topics of discussion) dimensions (Rowe & Snow, 2020). Conversational turns not only communicate linguistic and conceptual content, but they necessitate an interactive social exchange between the adult and child. Turn-taking exchanges may create a feedback loop in which the child has opportunities to practice their developing language with an attentive partner, while also providing the adult with increased opportunities to observe the child’s language level and adapt their speech to an optimal level of complexity. However, one limitation of the present methods is that LENA provides a window into the *frequency* of turn-taking, but does not provide information on the *content* of conversational exchanges, nor the *physical and social environments* in which they take place. Such information is critical to drive conclusions regarding the mechanisms by which conversational turns support children’s cognitive development. Future studies should investigate how the three dimensions of language input interact and evolve over ontogeny to support children’s language development and related neurocognitive domains.

Another limitation of this study is that the strongest results are indirect effects in absence of direct effects or correlations amongst concurrent change measures, which limits causal interpretations. This may also affect the interpretation about the direction of effects. For example, because the intervention also had a child component, it is possible that intervention-related changes in child language and brain morphometry drove conversational changes. However, we believe this explanation is unlikely, because the child-facing component of the intervention focused exclusively on attention and emotional regulation, and because there were no effects of the intervention on child vocalizations. The difference between the significant longitudinal correlations and the limited group effects on cognitive and brain measures appears to arise from large individual variability within both the intervention and control groups. Although the intervention did have significant effects on conversational turn-taking, about half of families exhibited increases and half decreases. It is unclear what factors drive within-family variation in communication practices, and how interventions act on this variation. It is striking, however, that a mere 10-20 weeks of fluctuating home language experience relates to fluctuations in language and cognitive abilities as well as brain anatomy. Such rapid cortical plasticity has been noted in other child and adolescent intervention studies—such as after 2 weeks of executive cognition training (Hoekzema et al., 2011), 6 weeks of reading intervention (Romeo, Christodoulou, et al., 2018), 5 weeks of inhibitory control training (Delalande et al., 2020), 12 weeks of exercise training (Szulc-Lerch et al., 2018), and 3 months of experience playing Tetris (Haier et al., 2009)—supporting the conclusion that rapid changes are possible and meaningful.

Additionally, the multifaceted nature of the intervention makes it difficult to pinpoint the mechanism underlying effects. While parent-child communication was a significant and pervasive part of the curriculum, the intervention involved many family practices designed to support child development, so it is possible that other factors may have resulted in changes both to turn-taking practices and children’s neurocognitive development. This is particularly likely for the effects of the intervention on executive functioning, which were mediated both by increased turn-taking and reductions in parents’ perceived stress. However, turn-taking changes were not correlated with changes in parent stress, parenting confidence, or discipline practices, making it unlikely that increased conversational turns is a mere proxy for other improvements to the parent-child relationship. Further research is need with targeted, communication-specific interventions to more precisely identify mechanisms linking language experience to neurocognitive development.

Additional limitations may constrain generalization. Because families were recruited from public charter schools which require entry to a lottery for enrollment, the sample may not be representative of broader populations of families who lack the resources to engage with such opportunities. Furthermore, the families who signed up for an intervention may have been more attuned to their children’s developmental needs. Additionally, families who had more barriers to attendance, such as inflexible work schedules or transportation uncertainty, may have been less likely to sign up to participate, attend assigned intervention sessions, and/or participate in the supplemental assessments. Future studies in population-based settings (e.g., Wong et al., 2020) are needed to ensure equal access to family support programs and reduce potential selection bias. Finally, this intervention was designed for families in Western societies where adult-child conversation, child expressiveness, and formal education are valued. As such, the neurocognitive effects of the intervention may not generalize to other cultures where such practices are rare or not valued, nor may such approaches be appropriate for widespread practice. It is critical that interventions to family practices take into account caregivers’ goals for their children’s development and futures.

This study is the first to reveal mechanisms of neuroplasticity that relate children’s language environments to their linguistic and cognitive development. Beyond illuminating neurobiological mechanisms, this study has important implications for education and public policy. It suggests the need for developing and expanding programs to support parents in providing high-quality language environments for their children. In parallel, it suggests potential benefits to reducing barriers that impede caregivers’ ability to spend time with children and engage in conversational turn-taking (Rowe, 2018; Ellwood-Lowe et al., 2020). Such multipronged approaches to family support may help to ensure optimal developmental outcomes for all children.

## Acknowledgements and Funding

Research was funded by the Walton Family Foundation (to M.R.W.), National Institute of Child Health and Human Development (F31HD086957 to RRR), Harvard Mind Brain Behavior Grant (to RRR), and a gift from David Pun Chan (to JDEG). We thank the University of Oregon Brain Development Lab (including Eric Pakulak, Zayra Longoria, Jimena Santillan, and Helen Neville) for providing the curriculum and training the interventionists to fidelity; Transforming Education, 1647 Families, and John Connolly; Glennys Sanchez, Marta Rivera, Samilla Quiroa, Barney Arnold, Violeta Vicario, and Blanca Valentin for delivering the intervention; the Athinoula A. Martinos Imaging Center at the McGovern Institute for Brain Research (MIT), including Atshusi Takahashi, Steve Shannon, and Sheeba Arnold for MRI assistance; Kelly Halverson, Emilia Motroni, Lauren Pesta, Veronica Wheaton, and Christina Yu for assistance in administering behavioral assessments; Tina Zhao, Sophia Diggs-Galligan, Jack Sandstedt, and Joshua Segaran for neuroimaging quality assurance; Maya L. Rosen for providing comments on an earlier version of this manuscript; and to all participating schools and families.

## Author Contributions

RRR and JDEG developed the study concept. RRR, JAL, ES, APM, MLR, MRW, and JDEG designed the study. RRR, JAL, STR, and MT collected the data. RRR, JAL, HMG, STR, and MT preprocessed, scored, and curated the data. RRR performed the data analysis and interpretation, under the supervision of JDEG and MLR. RRR drafted the manuscript and JDEG, MLR, APM, JAL, and MET provided revisions. All authors approved the final version of the manuscript for submission.

## References

Adolphs, R. (2009). The Social Brain: Neural Basis of Social Knowledge. Annual Review of Psychology, 60(1), 693–716.

APA Task Force on Race and Ethnicity Guidelines in Psychology. (2019). APA Guidelines on Race and Ethnicity: Promoting Responsiveness and Equity.

Brito, N. H., Troller-Renfree, S. V., Leon-Santos, A., Isler, J. R., Fifer, W. P., & Noble, K. G. (2020). Associations among the home language environment and neural activity during infancy. Developmental Cognitive Neuroscience, 43, 100780.

Bruce, J., McDermott, J. M., Fisher, P. A., & Fox, N. A. (2009). Using behavioral and electrophysiological measures to assess the effects of a preventive intervention: A preliminary study with preschool-aged foster children. Prev Sci, 10(2), 129–140.

Cohen, S., Kamarck, T., & Mermelstein, R. (1983). A global measure of perceived stress. Journal of Health and Social Behavior, 24(4), 385–396.

Cohen, S., & Williamson, G. (1988). Psychological stress in a probability sample of the united states. In S. Spacapan & S. Oskamp (Eds.), The social psychology of health: Claremont symposium on applied social psychology (pp. 31–67). Newbury Park, CA: Sage.

Delalande, L., Moyon, M., Tissier, C., Dorriere, V., Guillois, B., Mevell, K., … Borst, G. (2020). Complex and subtle structural changes in prefrontal cortex induced by inhibitory control training from childhood to adolescence. Developmental Science, 23(4), e12898.

DeMaster, D., Bick, J., Johnson, U., Montroy, J. J., Landry, S., & Duncan, A. F. (2019). Nurturing the preterm infant brain: Leveraging neuroplasticity to improve neurobehavioral outcomes. Pediatric Research, 85(2), 166–175.

Desikan, R. S., Segonne, F., Fischl, B., Quinn, B. T., Dickerson, B. C., Blacker, D., … Killiany, R. J. (2006). An automated labeling system for subdividing the human cerebral cortex on MRI scans into gyral based regions of interest. Neuroimage, 31(3), 968–980.

Diamond, A. (2013). Executive Functions. Annual Review of Psychology, 64(1), 135–168.

Duncan, G. J., & Magnuson, K. (2012). Socioeconomic status and cognitive functioning: Moving from correlation to causation. Wiley Interdisciplinary Reviews: Cognitive Science, 3(3), 377–386.

Dunn, L. M., & Dunn, D. M. (2007). Peabody picture vocabulary test (4th ed.). Bloomington, MN: Pearson.

Ellwood-Lowe, M. E., Foushee, R.; Srinivasan, M. (2020). What causes the word gap? Financial concerns may systematically suppress child-directed speech. PsyArXiv. https://doi.org/10.31234/osf.io/byp4k

Fedorenko, E., & Blank, I. A. (2020). Broca’s Area Is Not a Natural Kind. Trends in Cognitive Sciences, 24(4), 270–284.

Ferjan Ramirez, N., Lytle, S. R., Fish, M., & Kuhl, P. K. (2019). Parent coaching at 6 and 10 months improves language outcomes at 14 months: A randomized controlled trial. Developmental Science, 22, e12762.

Ferjan Ramírez, N., Lytle, S. R., & Kuhl, P. K. (2020). Parent coaching increases conversational turns and advances infant language development. Proceedings of the National Academy of Sciences, 117(7), 3484–3491.

Ferrer, E., O’Hare, E. D., & Bunge, S. A. (2009). Fluid reasoning and the developing brain. Frontiers in Neuroscience, 3(1), 46–51.

Fisher, P. A., Frenkel, T. I., Noll, L. K., Berry, M., Yockelson, M. (2016). Promoting healthy child development via a two-generation translational neuroscience framework: The filming interactions to nurture development video coaching program. Child Development Perspectives, 10(4), 251–256.

Ford, M., Baer, C. T., Xu, D., Yapanel, U., & Gray, S. (2008). The LENA language environment analysis system: Audio specifications of the DLP-0121: Technical report LTR-03-2.

Friederici, A. D. (2011). The brain basis of language processing: From structure to function. Physiological Review, 91(4), 1357–1392.

Gilkerson, J., Richards, J. A., & Topping, K. (2017). Evaluation of a LENA-based online intervention for parents of young children. Journal of Early Intervention, 39(4), 281–298.

Gilkerson, J., Richards, J. A., Warren, S. F., Montgomery, J. K., Greenwood, C. R., Kimbrough Oller, D., … Paul, T. D.. (2017). Mapping the early language environment using all-day recordings and automated analysis. American Journal of Speech Language Pathology, 26(2), 248–265.

Gilkerson, J., Richards, J. A., Warren, S. F., Oller, D. K., Russo, R., & Vohr, B. (2018). Language Experience in the Second Year of Life and Language Outcomes in Late Childhood. Pediatrics, 142(4).

Hagler, D. J., Jr., Saygin, A. P., & Sereno, M. I. (2006). Smoothing and cluster thresholding for cortical surface-based group analysis of fMRI data. Neuroimage, 33(4), 1093–1103.

Hagoort, P. (2014). Nodes and networks in the neural architecture for language: Broca’s region and beyond. Current Opinion in Neurobiology, 28, 136–141.

Haier, R. J., Karama, S., Leyba, L., & Jung, R. E. (2009). MRI assessment of cortical thickness and functional activity changes in adolescent girls following three months of practice on a visual-spatial task. BMC Research Notes, 2, 174.

Hayes, A. F. (2009). Beyond Baron and Kenny: Statistical mediation analysis in the new millennium. Communication Monographs, 76(4), 408–420.

Heidlage, J. K., Cunningham, J. E., Kaiser, A. P., Trivette, C. M., Barton, E. E., Frey, J. R., & Roberts, M. Y. (2020). The effects of parent-implemented language interventions on child linguistic outcomes: A meta-analysis. Early Childhood Research Quarterly, 50, 6–23.

Helms, J. E., Jernigan, M., & Mascher, J. (2005). The meaning of race in psychology and how to change it: a methodological perspective. American Psychologist, 60(1), 27–36.

Hirsh-Pasek, K., Adamson, L. B., Bakeman, R., Owen, M. T., Golinkoff, R. M., Pace, A., … Suma, K. (2015). The contribution of early communication quality to low-income children’s language success. Psychological Science, 26(7), 1071–1083.

Hoekzema, E., Carmona, S., Ramos-Quiroga, J. A., Barba, E., Bielsa, A., Tremols, V., … Vilarroya, O. (2011). Training-induced neuroanatomical plasticity in ADHD: a tensor-based morphometric study. Human Brain Mapping, 32(10), 1741–1749.

Hoff, E. (2003). The specificity of environmental influence: Socioeconomic status affects early vocabulary development via maternal speech. Child Development, 74(5), 1368–1378.

Huttenlocher, J., Vasilyeva, M., Cymerman, E., & Levine, S. (2002). Language input and child syntax. Cognitive Psychology, 45, 337–374.

Isbell, E., Stevens, C., Pakulak, E., Hampton Wray, A., Bell, T. A., & Neville, H. J. (2017). Neuroplasticity of selective attention: Research foundations and preliminary evidence for a gene by intervention interaction. Proc Natl Acad Sci U S A, 114(35), 9247–9254.

Klapwijk, E. T., van de Kamp, F., van der Meulen, M., Peters, S., & Wierenga, L. M. (2019). Qoala-T: A supervised-learning tool for quality control of FreeSurfer segmented MRI data. Neuroimage, 189, 116–129.

Kuhl, P. K. (2007). Is speech learning ‘gated’ by the social brain? Developmenta Science, 10(1), 110–120.

Leech, K. A., & Rowe, M. L. (2020). An intervention to increase conversational turns between parents and young children. Journal of Child Language, 1–14.

Leung, C. Y. Y., Hernandez, M. W., & Suskind, D. L. (2020). Enriching home language environment among families from low-SES backgrounds: A randomized controlled trial of a home visiting curriculum. Early Childhood Research Quarterly, 50, 24–35.

McGillion, M., Pine, J. M., Herbert, J. S., & Matthews, D. (2017). A randomised controlled trial to test the effect of promoting caregiver contingent talk on language development in infants from diverse socioeconomic status backgrounds. Journal of Child Psychology and Psychiatry, 58(10), 1122–1131.

Merz, E. C., Maskus, E. A., Melvin, S. A., He, X., & Noble, K. G. (2020). Socioeconomic disparities in language input are associated with children’s language-related brain structure and reading skills. Child Development, 91, 846–860.

Milgrom, J., Newnham, C., Anderson, P. J., Doyle, L. W., Gemmill, A. W., Lee, K., … Inder, T. (2010). Early sensitivity training for parents of preterm infants: Impact on the developing brain. Pediatr Res, 67(3), 330–335.

Miller, E. K., & Cohen, J. D. (2001). An integrative theory of prefrontal cortex function. Annual Review of Neuroscience, 24(1), 167–202.

Neville, H. J., Stevens, C., Pakulak, E., Bell, T. A., Fanning, J., Klein, S., & Isbell, E. (2013). Family-based training program improves brain function, cognition, and behavior in lower socioeconomic status preschoolers. Proceedings of the National Academy of Sciences of the United States of America, 110(29), 12138–12143.

Oberhuber, M., Hope, T. M. H., Seghier, M. L., Parker Jones, O., Prejawa, S., Green, D. W., & Price, C. J. (2016). Four Functionally Distinct Regions in the Left Supramarginal Gyrus Support Word Processing. Cerebral Cortex, 26(11), 4212–4226.

Pace, A., Luo, R., Hirsh-Pasek, K., & Golinkoff, R. M. (2017). Identifying pathways between socioeconomic status and language development. Annual Review of Linguistics, 3(1), 285–308.

Pierce, L. J., Reilly, E., & Nelson, C. A. (2020). Associations between maternal stress, early language behaviors, and infant electroencephalography during the first year of life. J Child Lang, 1–28.

Remer, J., Croteau-Chonka, E., Dean, D. C., 3rd, D’Arpino, S., Dirks, H., Whiley, D., & Deoni, S. C. L. (2017). Quantifying cortical development in typically developing toddlers and young children, 1-6 years of age. Neuroimage, 153, 246–261.

Reynolds, C. R. & Kamphaus, R. W., & (2015). BASC-3 Parenting relationship questionnaire. Minneapolis, MN: Pearson.

Richardson, H., Lisandrelli, G., Riobueno-Naylor, A., & Saxe, R. (2018). Development of the social brain from age three to twelve years. Nature Communications, 9(1), 1027.

Romeo, R. R., Christodoulou, J. A., Halverson, K. K., Murtagh, J., Cyr, A. B., Schimmel, C., … Gabrieli, J. D. E. (2018). Socioeconomic status and reading disability: Neuroanatomy and plasticity in response to intervention. Cerebral Cortex, 28(7), 2297–2312.

Romeo, R. R., Leonard, J. A., Robinson, S. T., West, M. R., Mackey, A. P., Rowe, M. L., & Gabrieli, J. D. E. (2018). Beyond the “30 million word gap:” Children’s conversational exposure is associated with language-related brain function. Psychological Science, 29(5), 700–710.

Romeo, R. R.*, Leonard, J. A.*, Scherer, E., Robinson, S. T., Takada, M., Mackey, A. P., … Gabrieli, J. D. E. (under review). Replication and extension of a family-based training program to improve cognitive abilities in young children. *Co-first authorship

Romeo, R. R., Segaran, J., Leonard, J. A., Robinson, S. T., West, M. R., Mackey, A. P., … Gabrieli, J. D. E. (2018). Language exposure relates to structural neural connectivity in childhood. Journal of Neuroscience, 38(36), 7870–7877.

Rowe, M. L. (2012). A longitudinal investigation of the role of quantity and quality of child-directed speech in vocabulary development. Child Dev, 83(5), 1762–1774.

Rowe, M. L. (2018). Understanding socioeconomic differences in parents’ speech to children. Child Development Perspectives, 12, 122–127.

Rowe, M. L., & Goldin-Meadow, S. (2009). Differences in early gesture explain SES disparities in child vocabulary size at school entry. Science, 323(5916), 951–953.

Rowe, M. L., & Snow, C. E. (2020). Analyzing input quality along three dimensions: interactive, linguistic, and conceptual. Journal of Child Language, 47(1), 5–21.

Rucker, D. D., Preacher, K. J., Tormala, Z. L., & Petty, R. E. (2011). Mediation analysis in social psychology: Current practices and new recommendations. Social and Personality Psychology Compass, 5(6), 359–371.

Schwab, J. F., & Lew-Williams, C. (2016). Language learning, socioeconomic status, and child-directed speech. Wiley Interdisciplinary Reviews: Cognitive Science, 7(4), 264–275.

Szulc-Lerch, K. U., Timmons, B. W., Bouffet, E., Laughlin, S., de Medeiros, C. B., Skocic, J., … Mabbott, D. J. (2018). Repairing the brain with physical exercise: Cortical thickness and brain volume increases in long-term pediatric brain tumor survivors in response to a structured exercise intervention. Neuroimage Clinical, 18, 972–985.

Tamis-LeMonda, C. S., Kuchirko, Y., & Song, L. (2014). Why is infant language learning facilitated by parental responsiveness? Curr Dir Psychol Sci, 23(2), 121–126.

Tettamanti, M., & Weniger, D. (2006). Broca’s area: A supramodal hierarchical processor? Cortex, 42(4), 491–494.

Tisdall, M. D., Hess, A. T., Reuter, M., Meintjes, E. M., Fischl, B., & van der Kouwe, A. J. (2012). Volumetric navigators for prospective motion correction and selective reacquisition in neuroanatomical MRI. Magnetic Resonance in Medicine, 68(2), 389–399.

van der Kouwe, A. J., Benner, T., Salat, D. H., & Fischl, B. (2008). Brain morphometry with multiecho MPRAGE. Neuroimage, 40(2), 559–569.

Walker, D., Greenwood, C., Hart, B., & Carta, J. (1994). Prediction of school outcomes based on early language production and socioeconomic factors. Child Development, 65, 606–621.

Wechsler, D. (2012). Wechsler preschool and primary scale of intelligence (4th ed.). Bloomington, MN: Pearson.

Weyandt, L. L., Clarkin, C. M., Holding, E. Z., May, S. E., Marraccini, M. E., Gudmundsdottir, B. G., … Thompson, L. (2020). Neuroplasticity in children and adolescents in response to treatment intervention: A systematic review of the literature. Clinical and Translational Neuroscience, 4(2).

Wiig, E. H., Semel, E. M., & Secord, W. (2013). Clinical evaluation of language fundamentals (5th ed.). Bloomington, MN: Pearson.

Wong, K., Thomas, C., & Boben, M. (2020). Providence talks: A citywide partnership to address early childhood language development. Studies in Educational Evaluation, 64, 100818.

Xu, D., Yapanel, U., & Gray, S. (2009). Reliability of the LENA language environment analysis system in young children’s natural home environment: Technical report LTR-05-2.

Zimmerman, F. J., Gilkerson, J., Richards, J. A., Christakis, D. A., Xu, D., Gray, S., & Yapanel, U. (2009). Teaching by listening: The importance of adult-child conversations to language development. Pediatrics, 124(1), 342–349.

