## Supplement for "Neuroplasticity associated with changes in conversational turn-taking following a family-based intervention"

Baseline correlations between SES, language environment, and child cognitive scores are reported in Supplementary Table 1.

**Supplementary Table 1. Correlations amongst baseline measures of SES, language environment, and cognition**

| | PE | FI | AW | CT | CV | Lang | NVIQ | EF |
|---|---|---|---|---|---|---|---|---|
| Parental Education (PE) | 1 | | | | | | | |
| Family Income (FI) | .638*** | 1 | | | | | | |
| Adult Words (AW) | .267† | .108 | 1 | | | | | |
| Conversational Turns (CT) | .331* | .259† | .772*** | 1 | | | | |
| Child Vocalizations (CV) | .353* | .287† | .481** | .819*** | 1 | | | |
| Language (Lang) | .368** | .384** | .409** | .427** | .376* | 1 | | |
| Nonverbal IQ (NVIQ) | .265† | .189 | .260† | .346* | .273† | .550*** | 1 | |
| Executive Functioning (EF) | .211 | .016 | .421** | .335* | .229 | .516*** | .374** | 1 |

†$p < 0.1$, *$p < 0.05$, **$p < 0.01$, ***$p < 0.001$

Matrix of zero-order correlations between SES measures, baseline LENA language environment measures, and baseline standardized cognitive assessments, collapsed across groups (measures taken before randomization). Because parental education and family income are ordinal, correlations with these variables are Spearman rank-order correlation coefficients ($\rho$), while all other coefficients are Pearson's product-moment correlation ($r$).

The main text reports effects of the intervention excluding three participants randomly assigned to the intervention group who completed all assessments but did not attend the intervention. In Supplementary Table 2, we report intent-to-treat analyses which include those three participants as they were assigned. By including these three participants, the effect of the intervention on conversational turns remained significant [$F(1,37)=5.18$, $p<.05$, partial $\eta^2=.12$; mean change intervention = +2.84; control = -7.01], and the effect of the intervention on adult words remained marginal [$F(1,37)=3.34$, $p=.076$, partial $\eta^2=.08$; mean change intervention = +216.40; control = -55.25]. The effect of the intervention on child vocalizations and composite language scores remained insignificant. The significant effect of the intervention on executive functioning was reduced to marginal significance [$F(1,50)=2.83$, $p=.099$, partial $\eta^2=.05$; mean change intervention = +.13; control = -.14], and the marginal effect of the intervention on nonverbal IQ was reduced to insignificance [$F(1,50)=2.03$, $p=.16$, partial $\eta^2=.04$; mean change intervention = +1.03; control = +.52]. The effect of the intervention on cortical thickness plasticity remained insignificant.

**Supplementary Table 2. Intent-to-Treat Analysis of Intervention Effects**

| Measure | Pre-Test | | Post-Test | | Time*Group Interaction $\eta^2$ ($p$) |
|---|---|---|---|---|---|
| | Intervention M (SD) [n] | Control M (SD) [n] | Intervention M (SD) [n] | Control M (SD) [n] | |
| Adult Words per Hour | 1145.95 (399.96) [25] | 1258.81 (489.72) [18] | 1392.20 (545.34) [23] | 1183.68 (451.35) [16] | .083 (.076)† |
| Conversational Turns per Hour | 47.49 (18.34) [25] | 55.77 (21.79) [18] | 50.70 (19.59) [23] | 48.71 (20.73) [16] | .123 (.029)* |
| Child Vocalizations per Hour | 204.09 (90.92) [25] | 222.84 (91.39) [18] | 209.12 (85.43) [23] | 233.59 (111.72) [16] | .000 (n.s.) |
| Language Composite | 98.44 (15.54) [27] | 100.18 (15.73) [25] | 101.69 (13.77) [27] | 103.88 (13.36) [25] | .001 (n.s.) |
| Nonverbal IQ Composite | 9.48 (2.10) [27] | 9.93 (2.23) [25] | 10.51 (2.12) [27] | 10.45 (2.26) [25] | .039 (.160) |
| Executive Functioning Composite | -.05 (.68) [27] | 0.05 (.77) [25] | 0.08 (.57) [27] | -.09 (.86) [25] | .054 (.099) † |

†$p < 0.1$, *$p < 0.05$

Descriptive and inferential statistics of standardized cognitive assessments and language environment measures, divided by assigned group and timepoint. Descriptive statistics include mean (M), standard deviation (SD), and subgroup sample size with count of valid measurements (n). Inferential statistics include effect size ($\eta^2$) and corresponding p-value derived from repeated measures ANOVAs with timepoint (pre-test or post-test) within-subjects and assigned group (intervention or control) between subjects. Three participants randomly assigned to the intervention did not attend any sessions (non-compliers) but are included here to examine intent-to-treat effects.
